# Changes in brain metabolite levels across childhood

**DOI:** 10.1101/2022.11.30.518618

**Authors:** Meaghan V. Perdue, Marilena M. DeMayo, Tiffany K. Bell, Elodie Boudes, Mercedes Bagshawe, Ashley D. Harris, Catherine Lebel

## Abstract

Metabolites play important roles in brain development and their levels change rapidly in the prenatal period and during infancy. Metabolite levels are thought to stabilize during childhood, but the development of neurochemistry across early-middle childhood remains understudied. We examined the developmental changes of key metabolites (total N-acetylaspartate, tNAA; total choline, tCho; total creatine, tCr; glutamate+glutamine, Glx; and myo-inositol, mI) using short echo-time magnetic resonance spectroscopy (MRS) in the anterior cingulate cortex (ACC) and the left temporo-parietal cortex (LTP) using a mixed cross-sectional/longitudinal design in children aged 2-11 years (ACC: N=101 children, 112 observations; LTP: N=95 children, 318 observations). We found age-related effects for all metabolites. tNAA increased with age in both regions, while tCho decreased with age in both regions. tCr increased with age in the LTP only, and mI decreased with age in the ACC only. Glx did not show linear age effects in either region, but a follow-up analysis in only participants with ≥3 datapoints in the LTP revealed a quadratic effect of age following an inverted U-shape. These substantial changes in neurochemistry throughout childhood likely underlie various processes of structural and functional brain development.

## Introduction

The preschool and early school years are marked by extensive cognitive, emotional, and social development as children gain independence, learn to interact with peers, and begin formal education. Substantial changes in brain structure and function occurs alongside this behavioral maturation (Long et al., 2017; Remer et al., 2017; Reynolds et al., 2019). Less is known about the development of the brain’s chemical make-up – specifically the metabolites that are involved in various aspects of brain development, metabolism, and neural signaling. By characterizing the changes in metabolites throughout childhood, we can understand the neurochemistry underpinning healthy brain development. Such insight may be useful to identify markers and mechanisms of disorders and disease, and to provide complementary information to our understanding of structural and functional development.

The primary brain metabolites measured with MRS include N-acetylaspartate, choline, creatine, glutamate/glutamine and myo-inositol, and each has a distinct developmental pattern (see Cichocka & Bereś, 2018 for a systematic review of metabolite develpoment across the lifespan). N-acetylaspartate (NAA) is primarily found in mature neurons and is thus considered a neuronal marker (Blüml et al., 2013; Ross & Sachdev, 2004). The interpretation of the NAA signal is complex, with several roles proposed for this metabolite, including neuronal metabolism, myelin lipid synthesis, synthesis of the neurotransmitter N-acetylaspartylglutamate (NAAG), and maintaining water balance by removing water from neurons (Hirrlinger & Nave, 2014; Moffett et al., 2007; Rae, 2014). NAA is often reported in combination with NAAG (labelled NAA+NAAG or total NAA [tNAA]) because the signals of these metabolites are not reliably separated with conventional MRS. Evidence from studies reporting water-referenced metabolite concentrations and those reporting metabolite ratios (NAA/Cr or NAA/Cho) converge to show rapid increases in tNAA from *in utero* through infancy (Blüml et al., 2013; Girard et al., 2006; Hashimoto et al., 1995; Kato et al., 1997; Kimura et al., 1995; Kok et al., 2002; Kreis et al., 1993, 2002; Patkee et al., 2021; van der Knaap et al., 1990; Vigneron, 2006), followed by gradual increases in childhood and adolescence (Blüml et al., 2013; Costa et al., 2002; Degnan et al., 2014; Hashimoto et al., 1995; Holmes et al., 2017; van der Knaap et al., 1990). Notably, there have also been reports of decrease (Bozgeyik et al., 2008; Devito et al., 2007) or stability (Giménez et al., 2004; Lam et al., 1998) of tNAA and NAA ratios (relative to Cr or Cho) in children/adolescents. These findings indicate possible region- or tissue-dependent effects and demonstrate the need for dense sampling across childhood along with the use of current methodological best-practices to characterize tNAA development.

Choline (Cho) is related to membrane turnover, including synthesis and repair of the myelin sheath (Blüml et al., 2013; Rae, 2014; Ross & Sachdev, 2004). The total choline (tCho) signal measured by MRS is primarily composed of phosphorylcholine and glycerophosphorylcholine, with a small contribution of Cho (Rae, 2014; Ross & Sachdev, 2004). tCho levels and Cho/Cr ratios are highest during the prenatal and neonatal stages, decrease rapidly during infancy, and are thought to stabilize in childhood (Blüml et al., 2013; Cady et al., 1996; Cichocka & Bereś, 2018; Degnan et al., 2014; Girard et al., 2006; Hashimoto et al., 1995; Holmes et al., 2017; Kimura et al., 1995; Kok et al., 2002; Kreis et al., 1993; Lam et al., 1998; van der Knaap et al., 1990). Patterns of change in tCho levels vary by tissue type, with more rapid early decreases in gray matter than white matter (Blüml et al., 2013). Myelination may largely account for the decreasing tCho levels during infancy, driven by the incorporation of phosphorylcholine into myelin sheath macromolecules (Blüml et al., 1999).

Creatine (Cr) contributes to the brain’s energy supply and is considered a marker of energy use (Blüml et al., 2013; Rackayova et al., 2017; Ross & Sachdev, 2004). The total creatine (tCr) signal consists of creatine and phosphocreatine. Developmental studies show a generally increasing trend of tCr levels during the prenatal period and infancy (Blüml et al., 2013; Girard et al., 2006; Kreis et al., 1993, 2002) which stabilizes by adolescence (Blüml et al., 2013; Degnan et al., 2014; Kreis et al., 1993). One longitudinal study showed increases in tCr from age 5-10 years, calling its putative stability into question and raising concern for its use as a reference metabolite, especially in pediatric samples (Holmes et al., 2017).

Glutamate is an excitatory neurotransmitter and serves metabolic functions in the Krebs cycle, the glutamate-glutamine cycle, nitrogen regulation, and formation of gamma-amino-butyric acid (GABA; Rae, 2014). Due to their overlapping peaks in the MRS spectrum, glutamate is often reported in combination with glutamine, which is an amino acid precursor of glutamate, GABA, and aspartate (Blüml et al., 2013; Ross & Sachdev, 2004). The combined glutamate + glutamine concentration is referred to as Glx. Findings regarding developmental changes in Glx levels are mixed. Several studies have reported increasing Glx in infancy (Blüml et al., 2013; Degnan et al., 2014; Kreis et al., 2002), and Blüml et al. (2013) found that Glx stabilized by two years of age. However, a longitudinal study of 5-10-year-old children points to more prolonged increases in Glx levels in both white matter and cortical and subcortical gray matter regions (Holmes et al., 2017). Studies with samples spanning childhood through young adulthood, including a recent large study (N=144) of adolescents and young adults, show age-related decreases in Glx and Glu/Cr ratios (Ghisleni et al., 2015; Perica et al., 2022; Raininko & Mattsson, 2010; Shimizu et al., 2017; Volk et al., 2019). An additional interesting finding from Perica and colleagues’ (2022) study revealed decreasing inter-subject variability in Glu/Cr with age. The authors posit that this change in variability could be driven by plasticity-related fluctuations in Glu/Cr during adolescence that stabilize in adulthood. Together, the evidence points to a complex, nonlinear trajectory of Glx development from infancy through young adulthood.

Myo-inositol (mI) is involved in cellular signaling and lipid synthesis, and is typically considered a glial marker (Blüml et al., 2013; Ross & Sachdev, 2004). Evidence converges to show that mI levels are highest *in utero* and rapidly decrease through infancy to stabilize by early childhood (Blüml et al., 2013; Cichocka & Bereś, 2018; Degnan et al., 2014; Girard et al., 2006; Kreis et al., 1993; Lam et al., 1998). The rapid decline in mI levels in the late prenatal-early postnatal period indicates the importance of this metabolite in brain development and suggests a likely role in myelination (Blüml et al., 2013).

As summarized above, prior MRS studies reveal distinct patterns of developmental change in brain metabolite levels, predominantly in the early postnatal months (Blüml et al., 2013; Kreis et al., 1993; Vigneron, 2006). These findings have led researchers to speculate that metabolite levels remain relatively stable across childhood and adolescence (Blüml et al., 2013; Lam et al., 1998). However, metabolite changes across childhood have been understudied, leaving a gap in the understanding of metabolic brain development. Few studies have examined metabolite changes across childhood, and those that have include small sample sizes with sparse data in the early childhood years, and most have examined metabolite ratios rather than absolute concentrations (Cichocka & Bereś, 2018). Furthermore, the existing data is primarily cross-sectional, meaning that developmental trajectories within individuals have not yet been characterized. Identifying patterns of neurochemical changes is an important step toward understanding the mechanisms that underlie the cognitive and behavioral changes during this crucial developmental period.

In this study, we used a mixed cross-sectional/longitudinal design to investigate changes in metabolite levels across childhood in the anterior cingulate cortex (ACC) and the left temporo-parietal cortex (LTP). These regions play important roles in cognitive development and have shown substantial structural (Norbom et al., 2020; Remer et al., 2017) and functional (Long et al., 2017; Xiao et al., 2016) changes in this age range. The ACC plays key roles in cognitive control and emotion regulation networks (Braver et al., 2021; Stevens et al., 2011), and the LTP is involved in attention, social cognition, and semantic processing (Numssen et al., 2021), and supports language and reading skills (Richlan, 2012). Both of these regions have been considered in prior MRS studies of childhood disorders (Horowitz-Kraus et al., 2018; Kossowski et al., 2019; Puts et al., 2020); characterizing development of metabolites in this region will provide important context for interpreting these and future studies. We measured concentrations of tNAA, tCho, tCr, Glx and mI using short echo MRS in 124 children ranging in age from 2-11 years (analyses include 430 total observations across two datasets [ACC: N=112 observations; LTP: N=318 observations). We predicted that tNAA and tCr levels would increase across the age range, tCho and mI would decrease to stabilize in middle childhood, and Glx would increase slightly or show no clear developmental change. We expected similar patterns of change in both regions of interest. The large sample size and substantial longitudinal data within the LTP dataset afforded the ability to conduct additional analyses in a subset of participants with data at 3 or more time points (51 individuals, N=250 observations) to test for nonlinear patterns of metabolite development and identify ages at which metabolite levels reach minimum or maximum values.

## Methods

### Participants and data collection timeline

Data was drawn from an accelerated longitudinal study of brain development across early-middle childhood conducted in Calgary, AB, Canada (Reynolds et al., 2020). None of the participants had diagnosed neurological, genetic, or neurodevelopmental disorders, and all were born full term (≥37 weeks gestation). Participants were invited to return for MRI scans semi-annually between ages 2-4 years and annually thereafter.

#### Ethics statement

Parents provided written informed consent and children provided verbal assent. This study was approved by the conjoint health research ethics board (CHREB) at the University of Calgary (REB13-0020).

The present analyses include 430 total datasets from 124 participants (61 female, mean age = 5.46, SD = 1.93, range 2.34-11.13 y). These data were drawn from a database of 598 scans completed as of November 2021 and were selected based on completion of the MRS sequence and subsequent quality assurance. 130 of the 598 total scans were excluded because no MRS data was collected (due to subjects wanting to get out of the scanner, time constraints, and/or poor quality of T1-weighted scans preventing accurate voxel placement); 1 scan was excluded due to MRS acquisition outside of the ACC or LTP (voxel acquired in inferior frontal gyrus during study development); 20 scans were excluded because the MRS data files could not be retrieved; an additional 17 scans were excluded during the quality checking procedures, detailed in the following sections). Median household income for the sample was $150,000 - $174,999 CAD (range: <$25,000 – >$175,000), median level of maternal education was undergraduate degree (range: completed high school - postgraduate degree), and median level of paternal education was undergraduate degree (range: some high school - postgraduate degree). The full sample includes two analysis subsets: (1) data acquired from the midline ACC (N=112 datasets, 101 participants [47 females], mean age = 4.09 years, SD = 1.11, age range: 2.34 - 7.36 years) and (2) data acquired from the LTP region in an overlapping set of participants (N=318 datasets, 95 participants [47 female], mean age = 5.96 years, SD = 1.92, age range: 2.41 - 11.13 years). MRS scans were acquired from the ACC at visit 1 (though ACC data was acquired at multiple time points for five subjects) and from the LTP at follow-up visits for most participants, with up to nine longitudinal data points (Figure 1). This resulted in a primarily cross-sectional data set for the ACC and a longitudinal dataset for the LTP.

**Figure 1.**
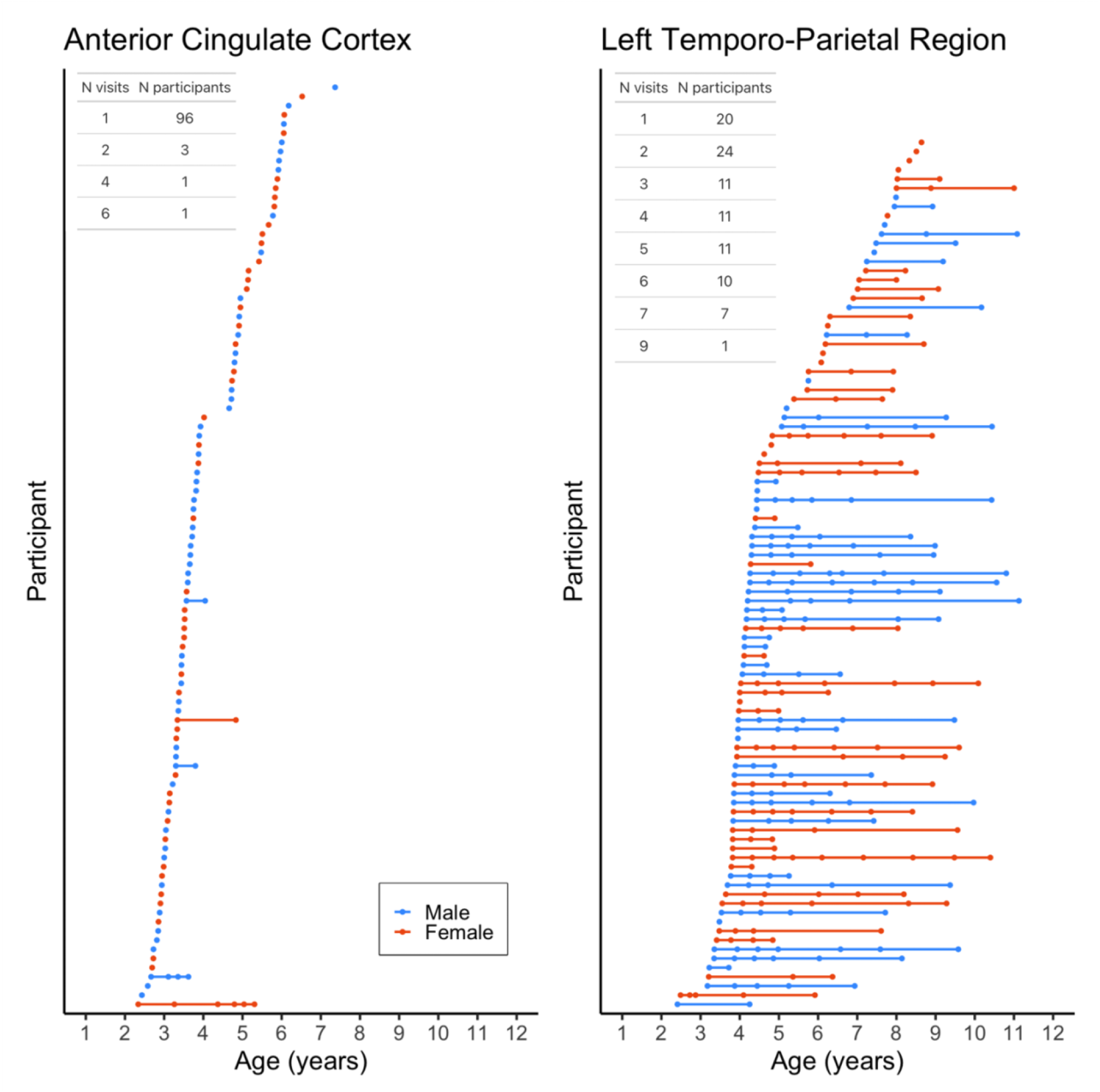
Age at scan acquisition for the ACC (left) and LTP (right). Each scan is represented by a circle; each participant is shown in a different row with their scans connected by a straight line. Inset tables indicate the number of participants who completed each number of visits.

### Data Acquisition

Prior to participation, families were provided with training materials to familiarize children with MRI and were offered an opportunity to complete a mock-MRI training session (see Reynolds, et al. 2020 for details). Our lab has demonstrated high success rates in MRI scanning of young children with and without mock-MRI training (Thieba et al., 2018).

MRI sessions were conducted using a research-dedicated scanner at the Alberta Children’s Hospital (Calgary, AB, Canada) by staff who are highly skilled in pediatric neuroimaging. Anatomical images and metabolite data were acquired using a 3T GE MR750w MR system with a 32-channel head coil. Children were scanned while watching a movie or during natural sleep; no sedation was used. T1-weighted anatomical images were acquired using a spoiled gradient echo sequence (210 axial slices; 0.9 x 0.9 x 0.9mm resolution, TR = 8.23 ms, TE = 3.76 ms, flip angle = 12°, matrix size = 512 x 512, inversion time = 540 ms). These images were reformatted to provide axial, sagittal, and coronal views at the scanner which were used for placement of spectroscopy voxels. MRS data were acquired using short echo time Point RESolved spectroscopy (PRESS; TE = 30 ms, TR = 2000 ms, 96 averages, 20 x 20 x 15 mm voxels). A strong body of literature demonstrates the test-retest reliability and stability of PRESS sequences and supports the validity of MRS measurements for longitudinal research (Baeshen et al., 2020; Fayed et al., 2009; Gasparovic et al., 2011; Soreni et al., 2010; Volk et al., 2018, 2019). MRS voxels were placed in either the anterior cingulate cortex (ACC) or in the left temporal-parietal area (LTP) by trained members of the research team according to detailed instructions and reference images. The ACC voxel was localized anterior to and at approximately the same level as the genu of the corpus callosum viewed on a midsagittal slice and consisted almost entirely of gray matter (Figure 2). The LTP voxel was localized to capture the left angular gyrus based on all three image planes, and included portions of the supramarginal gyrus, parietal operculum, and posterior superior temporal gyrus due to the extent of the voxel (Figure 2). The LTP voxel consisted of gray and white matter, excluding CSF to the extent possible.

**Figure 2.**
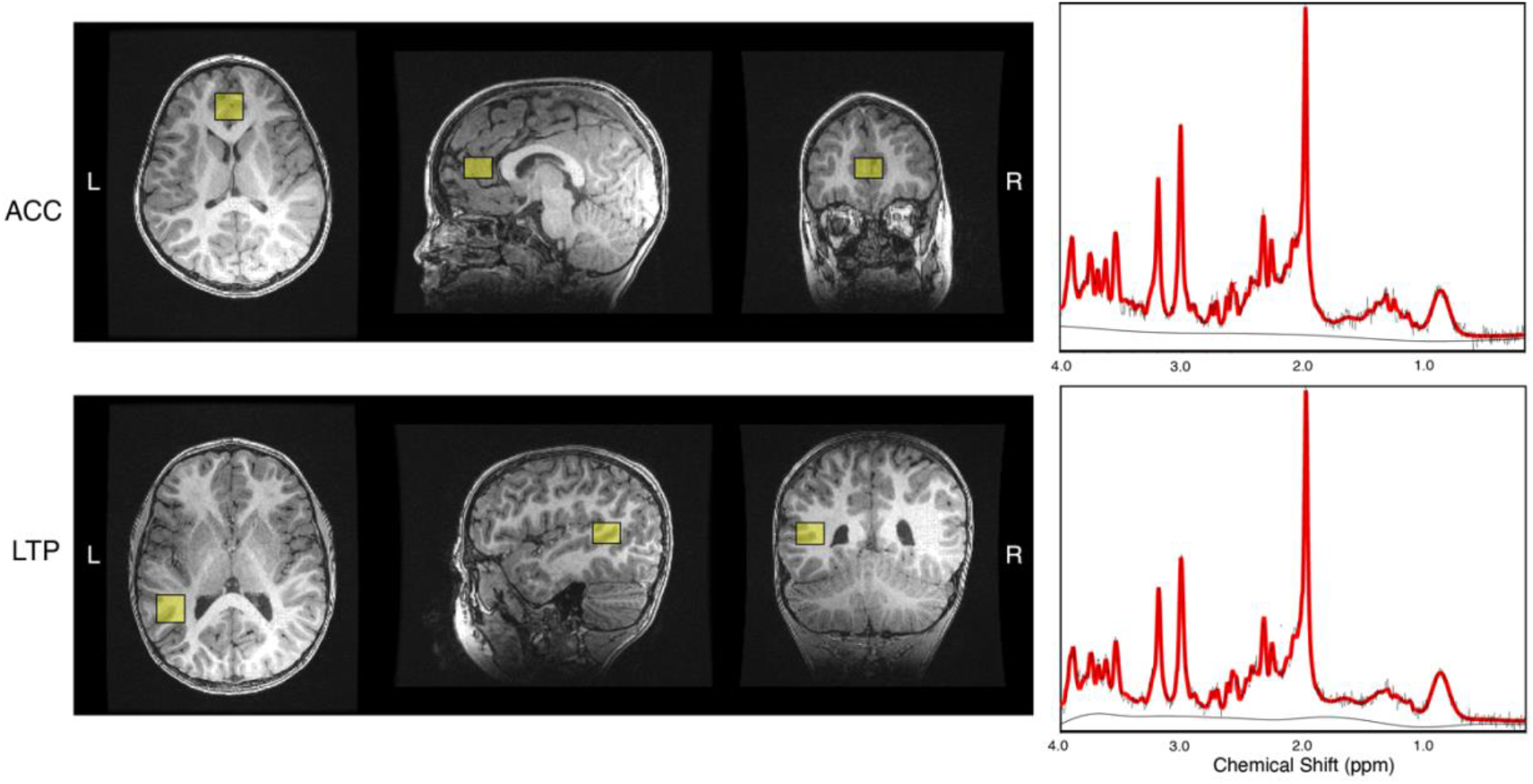
Example voxel placement and spectra (raw data shown in black; spectra fitted with LCModel shown in red) for the anterior cingulate cortex (ACC; top) and left temporo-parietal (LTP; bottom).

This data was acquired as part of a larger neuroimaging study including multiple sequences. MRS data were acquired in the second half of the protocol, and ∼20% of cases (130 of 598 datasets) stopped scanning prior to the MRS sequence due to time constraints and/or excessive motion.

### MRS data processing and analysis

The PRESS acquisition was pre-processed with the FID-A toolbox (Simpson et al., 2017). Pre-processing included coil combination, removal of bad averages, frequency alignment, and phase correction, according to recent recommendations (Near et al., 2020). Metabolites were then fit using LCModel v6.3 (Provencher, 1993). The metabolites of interest were: tNAA, tCr, tCho, mI and Glx. The basis set used for quantification included alanine, ascorbate, aspartate, choline, citrate, creatine, ethanol, GABA, glycerophosphocholine, glutathione, glucose, glycine, glutamine, glutamate, water, mI, lactate, NAA, NAAG, phosphocholine, phosphocreatine, phosphoryl ethanolamine, scyllo-inositol, taurine and beta-Hydroxybutyrate, simulated using the FID-A toolbox (Simpson et al., 2017) with sequence specific timings and RF pulse shapes. Fitted spectra were visually inspected for quality assurance and datasets with poor spectra were excluded from analysis (ACC: n=5; LTP: n=6). Quantitative quality metrics (signal-to-noise ratio [SNR] and linewidth [LW] for NAA) were obtained using the “op_getLW.m” and “op_getSNR.m” functions in FID-A. The Gannet CoRegStandAlone function (Harris et al., 2015), which calls spm12 segmentation (Ashburner & Friston, 2005), was used to quantify tissue content within the MRS voxels by co-registering them to the T1-weighted image acquired during the same scanning session. Co-registration and segmentation outputs were visually inspected to ensure accurate localization and segmentation of the MRS voxels. Datasets in which MRS voxels were not accurately located in the ACC or LTP were excluded from analysis (ACC: n = 3; LTP: n = 1). Absolute metabolite values are expressed in molal units (moles/kg), as per the approach recommended in the MRS consesnsus statement on metabolite quantification (Near et al 2020) and also described by Gasparovic et al (2006). This method accounts for the differential water T1- and T2-relaxation in white matter, gray matter and CSF in the voxel to enable absolute metabolite measurements. The scripts for the correction were developed by (DeMayo et al., 2023), and are publicly available (https://github.com/HarrisBrainLab/TissueCorrections). However, this approach does not address differences in metabolite concentrations between white matter and gray matter (as suggested in (Gasparovic et al., 2018)) as this ratio is not agreed upon for all metabolites (and may change with development). For this reason, the gray matter tissue fraction was included as a covariate in analyses.

The gray matter tissue fraction of each MRS voxel was calculated for each participant as the fraction of voxel volume composed of gray matter divided by total tissue within the voxel (sum of the fractions of voxel volume for gray matter (GM) and white matter (WM; formula: fGM/[fGM + fWM]). This tissue fraction was applied as a fixed effect in subsequent analyses to account for individual differences in tissue composition of the voxels to ensure that the effects observed in our models are driven by age and not tissue content of the voxels. The gray matter tissue fraction was maintained in final models only if including tissue fraction significantly improved model fit. Our model-fitting approach is detailed in the following section.

### Data analysis

Prior to analysis, distributions of metabolite values were checked for outliers by visually inspecting box plots and performing the Rosner’s Test for Outliers with the EnvStats R package (Millard, 2013). In addition to the previous quality assurance, spectra of datasets identified as outliers were visually inspected and excluded from analysis if artifacts were apparent around the peak value for the given metabolite and/or Cramer-Rao lower bounds estimates exceeded 20% (ACC: 1 dataset excluded from analysis of mI and tCho, 1 excluded from mI only; LTP: 2 datasets excluded from all analyses, 1 excluded from tNAA, 1 excluded from Glx, 4 excluded from mI). The distributions were visually checked using density plots before and after removal of outliers to confirm normal distribution of the data, and that the data meet the normality assumption required for Rosner’s Test.

Relationships between metabolite concentrations and age were investigated using linear mixed-effects modelling with the lme4 (Bates et al., 2015) and lmerTest (Kuznetsova et al., 2017) packages in RStudio version 2021.9.0.351 (RStudio Team, 2020). Datasets from the ACC and LTP voxels were analyzed independently, and separate models were fitted for each metabolite. First, linear growth models were defined with metabolite concentration as the dependent variable, age as a fixed effect, and participant as a random effect to account for repeated measures using the formula: metabolite ∼ age + (1 | participant). GM tissue fraction, sex, and the interaction between age and sex were sequentially added to the linear growth models and compared to the previous model to identify the best-fitting model for each metabolite. The best-fitting models were selected based on the likelihood ratio test criterion with alpha set to .05, maintaining the more parsimonious model when the likelihood ratio test did not show significant differences in model fit. False Discovery Rate (FDR) correction was performed within each region (ACC and LTP) to correct for running five models (Benjamini & Hochberg, 1995). Both corrected and uncorrected results are reported.

### Longitudinal analysis

To test for nonlinear effects of age, we conducted additional analysis of participants in the LTP sample who had MRS data for ≥3 time points. This analysis included 250 datasets from 51 subjects (23 females, mean age = 5.91 years, SD = 1.91, age range: 2.49-11.13). Based on model fitting for the full sample, we first fit mixed-effects models for each metabolite with metabolite concentration as the dependent variable, age and GM tissue fraction as fixed effects, and participant as a random effect using the formula: metabolite ∼ age + GM tissue fraction + (1 | participant). A quadratic age term was then added (formula: metabolite ∼ age + GM tissue fraction + age^2^ + (1 | participant)). The best-fitting model for each metabolite was selected based on the likelihood ratio test criterion with alpha set to .05. Multiple comparisons for running 5 models were corrected within each factor (e.g., age) using FDR (Benjamini & Hochberg, 1995). Both corrected and uncorrected *p*-values are reported.

### Post-hoc analysis of quality metrics

We conducted follow-up analyses to tests relationships between age and quality metrics (SNR and LW) using mixed-effects models. Quality metrics that were significantly associated with age were then included in post-hoc analysis of age-metabolite relationships to test whether the effects were driven by data quality.

#### Data Availability

De-identified participant demographics, metabolite concentration values, and analysis code are available at https://osf.io/q4t8j/?view_only=98430a2682b14be6a946a3984c31436f. T1-weighted anatomical images are available through the Calgary preschool magnetic resonance imaging dataset repository: https://osf.io/axz5r/ (Reynolds et al., 2020).

## Results

### Sample characteristics

Mean metabolite concentrations, GM tissue fractions, and quality metrics for each MRS voxel are reported in Table 1.

**Table 1.** Descriptive statistics for concentrations of metabolites (mmol/kg), Cramer-Rao Lower Bounds (CRLBs), gray matter tissue fraction (fGM/[fGM+fWM]), linewidth and signal-to-noise ratio (SNR) in each voxel. “N obs.” indicates total number of observations including longitudinal data within participants. Linewidth (FWHM) and SNR are reported for NAA.

|  | ACC |  |  | LTP |  |  |
| --- | --- | --- | --- | --- | --- | --- |
|  | N obs. | Mean | SD | N obs. | Mean | SD |
| tNAA (mmol/kg) | 112 | 15.66 | 1.26 | 318 | 14.98 | 1.03 |
| tCho (mmol/kg) | 111 | 2.60 | 0.34 | 318 | 2.29 | 0.27 |
| tCr (mmol/kg) | 112 | 12.72 | 1.03 | 318 | 10.82 | 0.91 |
| Glx (mmol/kg) | 112 | 26.84 | 2.58 | 317 | 21.53 | 2.16 |
| ml (mmol/kg) | 110 | 7.7 | 1.52 | 314 | 6.38 | 1.18 |
| tNAA (CRLB) | 112 | 2.64 | 0.60 | 318 | 2.20 | 0.47 |
| tCho (CRLB) | 111 | 4.60 | 1.28 | 318 | 4.16 | 0.82 |
| tCr (CRLB) | 112 | 2.81 | 0.56 | 318 | 2.54 | 0.55 |
| Glx (CRLB) | 112 | 4.82 | 1.04 | 317 | 4.95 | 0.80 |
| ml (CRLB) | 110 | 12.28 | 5.36 | 314 | 11.51 | 4.67 |
| GM Tissue Fraction | 112 | 0.96 | 0.03 | 318 | 0.64 | 0.11 |
| linewidth (Hz) | 112 | 7.38 | 3.17 | 318 | 6.67 | 2.22 |
| SNR | 112 | 57.29 | 10.64 | 318 | 71.48 | 13.82 |

### Age effects in the anterior cingulate cortex (ACC)

The best fitting model for all metabolites measured in the ACC was defined by the formula: metabolite ∼ age + (1 | participant). GM tissue fraction and sex had no significant effects and did not improve model fit, so these variables were not included in the final models.

tNAA concentration increased with age (*beta* = 0.25, *p* = .016) and tCho decreased with age (*beta* = -0.07, *p* = .018). A trending decrease was found for mI (*beta* = -.22, *p* = .083), but this effect was not significant. The models did not show significant age effects for tCr or Glx. Results are summarized in Table 2 and depicted in Figure 3.

**Figure 3.**
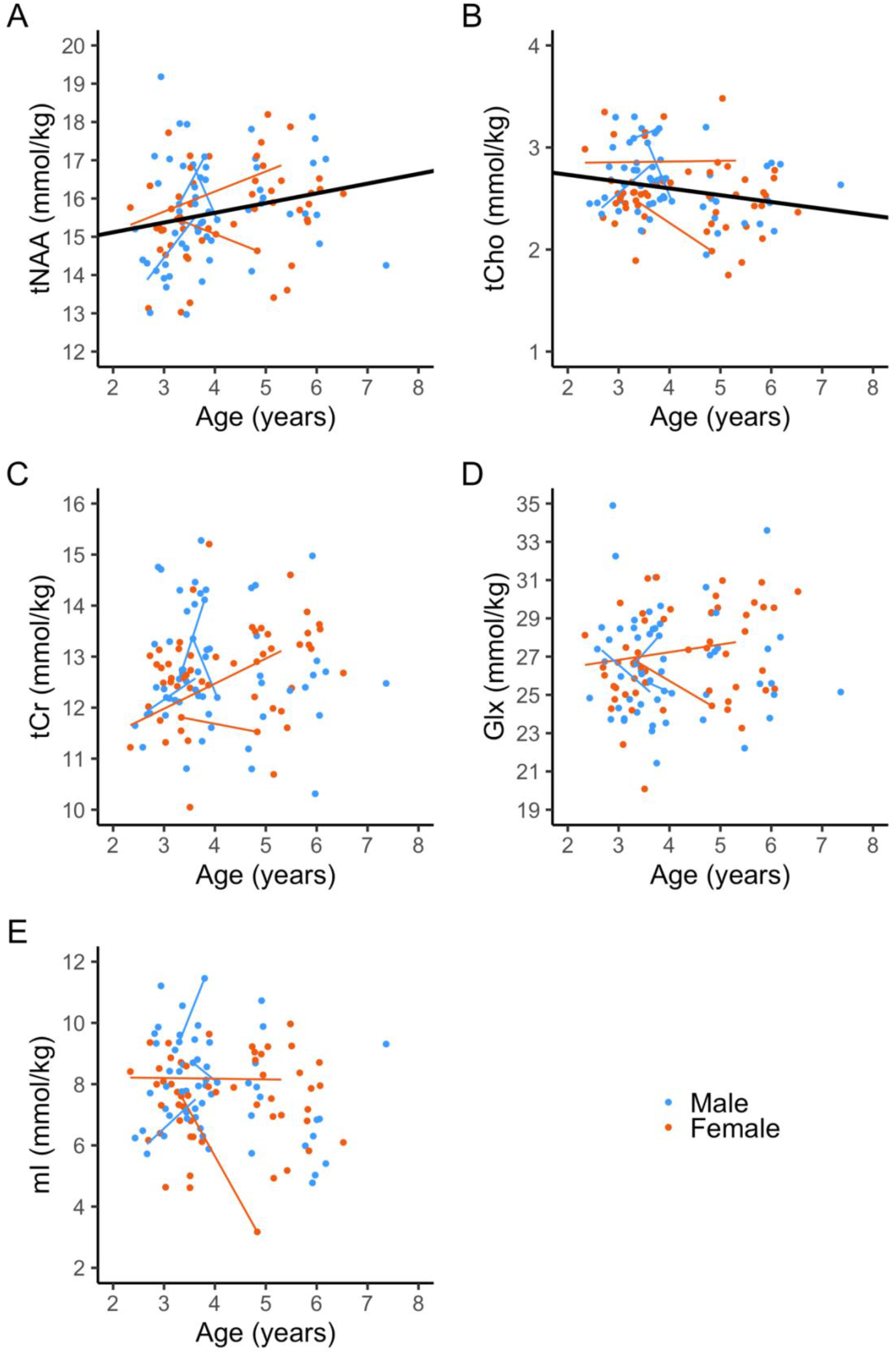
Relationships between age and metabolite concentrations (mmol/kg) in the ACC. Model fit lines shown in black for significant age effects only; individual participants’ fit lines shown in red and blue for participants with multiple scans.

**Table 2.** Summary of age effects in the ACC. Significant age effects (corrected for multiple comparisons) are presented in bold. Total percent change for significant models based on predicted values at ages 2 and 8 years.

| Metabolite | N obs. | Age effect |  |  |  |  | % change predicted |
| --- | --- | --- | --- | --- | --- | --- | --- |
| | | Intercept | beta [95% CI] | Std. beta [95% CI] | $p_{unc.}$ | $p_{FDR}$ | |
| tNAA | 112 | **14.61** | **.25 [.05, .46]** | **.23 [.04, .41]** | **.016** | **.045** | **10.10** |
| tCho | 111 | **2.87** | **-.07 [-.12, -.01]** | **-.22 [-.4, -.04]** | **.018** | **.045** | **-14.67** |
| tCr | 112 | 12.31 | .10 [-.07, .27] | .11 [-.07, .3] | .239 | .254 | NA |
| Glx | 112 | 25.82 | .25 [-.18, .68] | .11 [-.08, .29] | .254 | .254 | NA |
| ml | 110 | 8.61 | -.22[-.47, .03] | -.16 [-.35, .02] | .083 | .138 | NA |

### Age effects in the left-temporo-parietal region (LTP)

The best-fitting model for all metabolites in the LTP included age and GM tissue fraction as fixed effects, and was defined by the formula: metabolite ∼ age + tissue fraction + (1 | participant). Sex did not show any significant effects and did not improve fit of any models.

Our models showed age-related increases in tNAA (*beta* = 0.18, *p* < .001) and tCr (*beta* = .05, *p* = .025). tCho concentration decreased with age (*beta* = -.05, *p* <.001). Age was not significantly associated with mI or Glx. Results are summarized in Table 3 and depicted in Figure 4.

**Figure 4.**
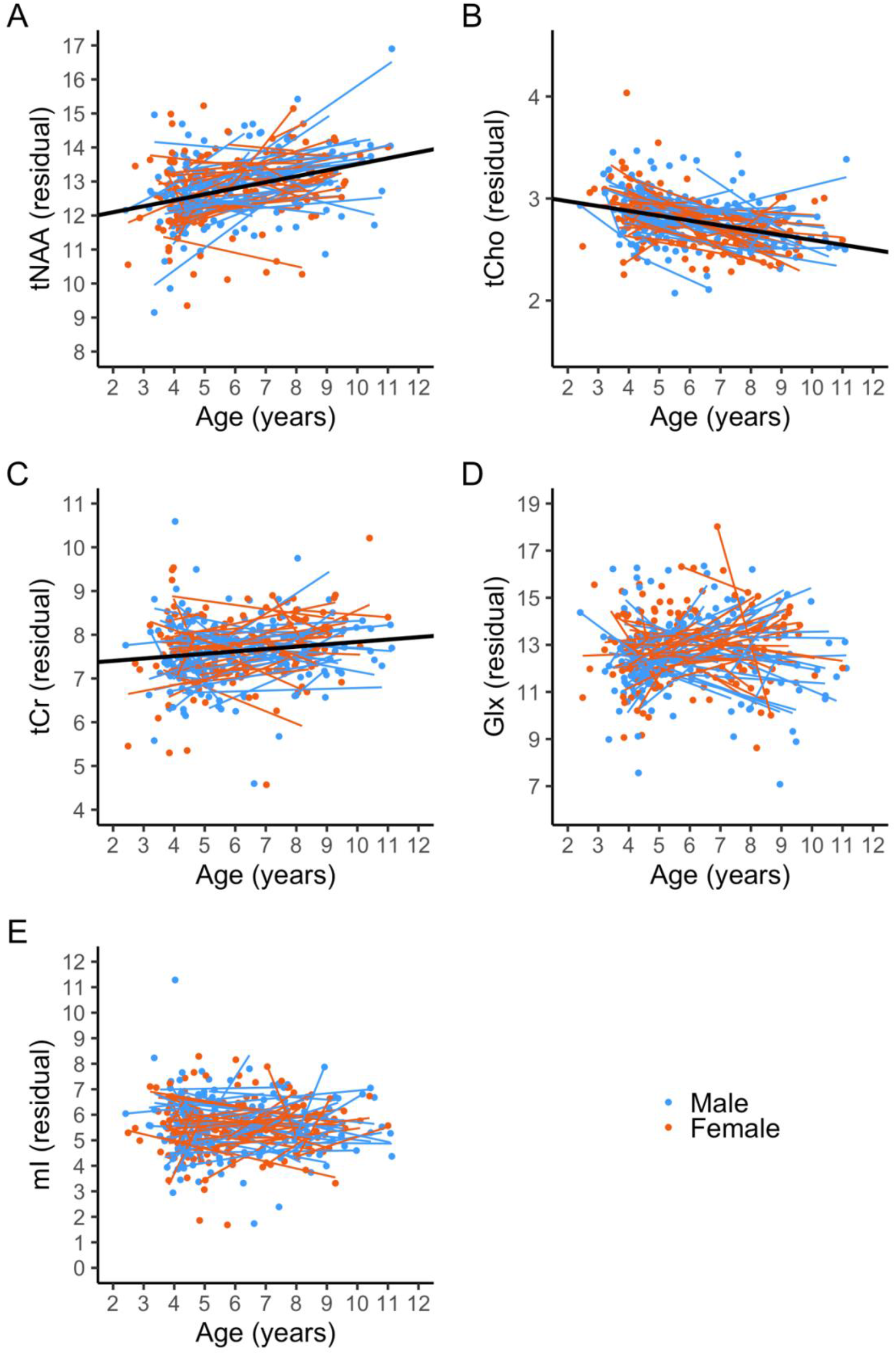
Relationships between age and metabolite concentrations in the LTP, controlling for tissue fraction. Model fit lines shown in black for significant age effects only; individual participants’ fit lines shown in red and blue for participants with multiple scans.

**Table 3.** Summary of age effects in the LTP. Significant age effects (corrected for multiple comparisons) are presented in bold. Total percent change for significant models based on predicted values at ages 2 and 11 years.

| Metabolite | N obs. | Age effect |  |  |  |  | % change predicted |
| --- | --- | --- | --- | --- | --- | --- | --- |
| | | Intercept | $\beta$ [95% CI] | Std. $\beta$ [95% CI] | $p_{unc.}$ | $p_{FDR}$ | |
| tNAA | 317 | **11.74** | **.18[.12, .24]** | **.33[.22,.43]** | **<.001** | **<.001** | **13.13** |
| tCho | 318 | **3.07** | **-.05[-.06, -.03]** | **-.34[-.45, -.23]** | **<.001** | **<.001** | **-14.37** |
| tCr | 318 | **7.29** | **.05[.01, .10]** | **.11[.01, .21]** | **.025** | **.042** | **6.60** |
| Glx | 317 | 12.49 | .04[-.07, .14] | .03[-.06, .12] | .497 | .621 | NA |
| ml | 314 | 5.58 | 0[-.08, .07] | 0[-.12, .12] | .958 | .958 | NA |

### Post-hoc analysis of quality metrics

We tested relationships between age and quality metrics (linewidth and SNR of the NAA peak) and found that SNR increased with age in the LTP dataset only (*beta* = 2.13, *p* < .001; Supplementary Table S1 & Figure S1); no significant age effects were found for linewidth in either voxel (Supplementary Tables S1 & S2). Thus, SNR of the NAA peak was added to the models testing age-metabolite relationships in the LTP. Age effects for tNAA and tCho remained consistent with the main analyses (tNAA: *beta* = .16, *p* < .001; tCho: *beta* = -.04, *p* < .001), however the tCr effect was no longer significant (*beta* = .04, *p* = .118). Statistics for metabolite-age effects for models including SNR as a covariate are reported in Supplementary Table S3.

### Analysis of nonlinear age effects in participants with LTP data from 3 or more time points

We conducted a follow-up analysis in a sub-set of participants who completed ≥3 scans in the LTP (n=51 subjects; total datasets = 250) to test for nonlinear relationships between age and metabolite concentrations. We first tested the linear age models controlling for tissue fraction in this longitudinal sub-sample, then added a quadratic age term to see if it significantly improved model fit.

**Table 4.** Mean concentrations of metabolites (mmol/kg) and mean gray matter tissue fraction (GM/[GM+WM]) in the participants with ≥3 LTP datasets. “N obs.” indicates total number of observations accounting for longitudinal data within participants.

|  | N obs. | Mean | SD |
| --- | --- | --- | --- |
| tNAA | 249 | 14.97 | 1.08 |
| tCho | 250 | 2.29 | 0.28 |
| tCr | 250 | 10.84 | 0.95 |
| Glx | 249 | 21.62 | 2.18 |
| ml | 247 | 6.40 | 1.20 |
| GM Tissue Fraction | 250 | 0.65 | 0.11 |

For tNAA and tCr, the quadratic age term did not improve model fit, and the linear age effects were significant and positive (tNAA: *beta* = .18, *p* <.001; tCr: *beta* = .07, *p* =.011), consistent with the effects observed in the full sample. The quadratic age term significantly improved model fit for tCho (*beta*_age_ = -.19, *p*_age_ <.001; *beta*_age^
2_ = .01, *p*_age^
2_ = .011) and Glx (*beta*_age_ = 1.17, *p*_age_ =.001; *beta*_age^
2_ = -.08, *p*_age^
2_ = .003). tCho decreased in early childhood, then slowed and hit a predicted minimum at age 8.8 years. Glx showed a modest increase across early childhood to peak at age 6.9 years followed by a decrease. No significant linear or quadratic effects of age were found for mI, consistent with the full LTP sample. Results are summarized in Table 5 and depicted in Figure 5.

**Figure 5.**
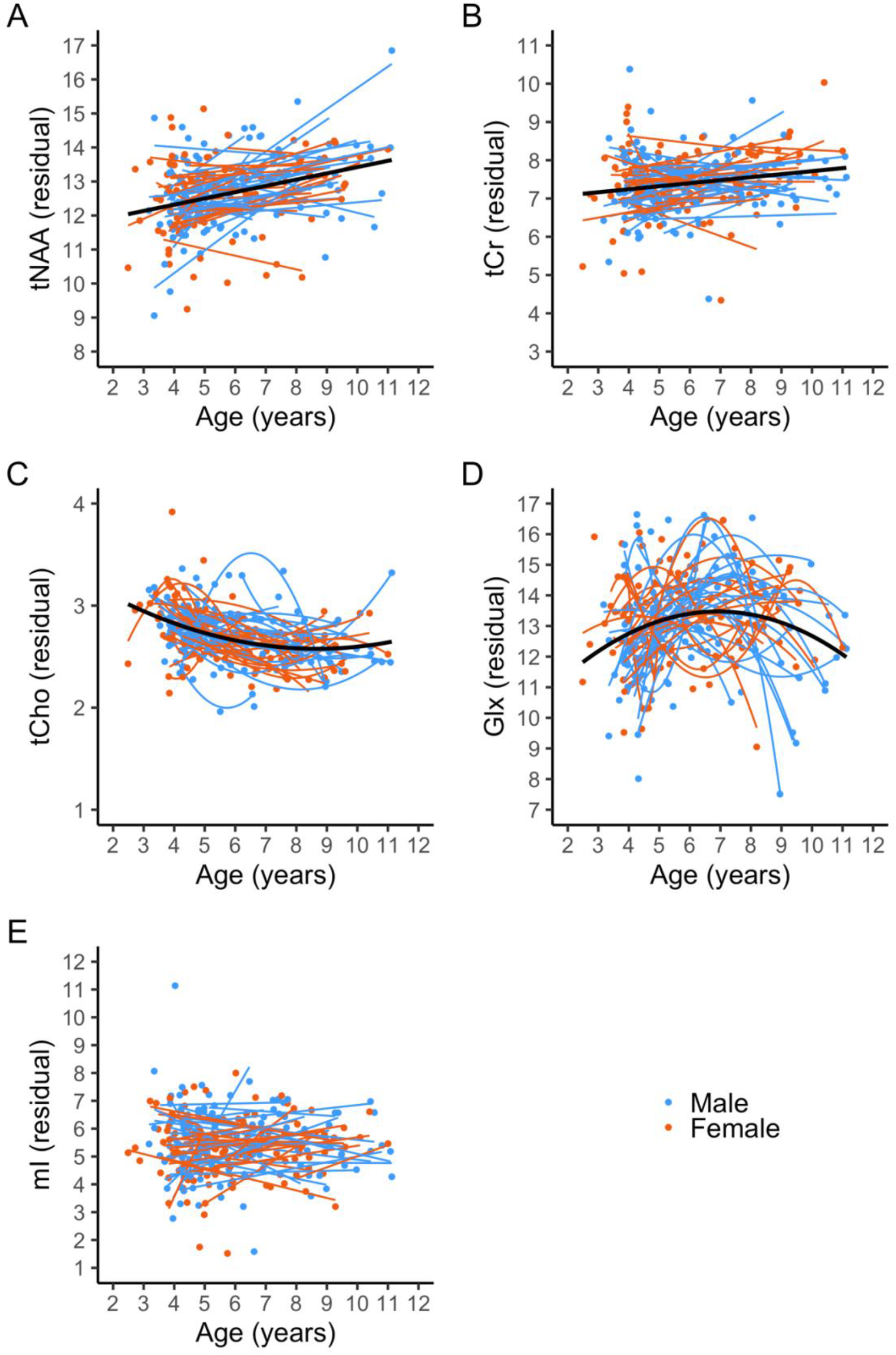
Linear and quadratic relationships between age and metabolite concentrations in the LTP sub-sample with ≥3 scans. Model fit lines shown in black for significant effects only; individual participants’ fit lines shown in red and blue.

**Table 5.** Summary of linear and quadratic age effects in sub-sample of participants with ≥3 scans in the LTP. Significant age and age^2^ effects (corrected for multiple comparisons) are presented in bold.

| Metabolite | N obs. | Intercept | Age effect |  |  |  | Quadratic age effect |  |  |  |
| --- | --- | --- | --- | --- | --- | --- | --- | --- | --- | --- |
| | | | beta[95% CI] | Std. Beta[95% CI] | $p_{unc.}$ | $p_{FDR}$ | beta[95% CI] | Std. Beta[95% CI] | $p_{unc.}$ | $p_{FDR}$ |
| tNAA | 249 | 11.62 | **0.18[.11, .25]** | **.32[.19, .44]** | **<.001** | **<.001** | NA | NA | NA | NA |
| tCho | 250 | 3.41 | **-.19[-.29, -.09]** | **-1.31[-2.03, -.59]** | **<.001** | **<.001** | **.01[.00, .02]** | **1.0[.27, 1.73]** | **.008** | **.011** |
| tCr | 250 | 6.97 | **.07[.02, .13]** | **.15[.03, .26]** | **.011** | **.01** | NA | NA | NA | NA |
| Glx | 249 | 9.43 | **1.17[.48, 1.86]** | **1.02[.41, 1.62]** | **.001** | **.001** | **-.08[-.14, -.03]** | **-.98[-1.6, -.37]** | **.002** | **.003** |
| ml | 247 | 5.46 | 0[-.09, .08] | 0[-.14, .13] | .955 | .955 | NA | NA | NA | NA |

## Discussion

Here, we show that multiple brain metabolites (tNAA, tCho, tCr, and Glx) continue to change substantially across early-middle childhood (age 2-11 years), indicating that maturation continues after rapid development in infancy (Blüml et al., 2013). These changes likely underlie ongoing structural and functional brain development during this age range.

tNAA concentrations increased with age throughout childhood. This finding corroborates evidence from cross-sectional studies (Blüml et al., 2013; Costa et al., 2002; Degnan et al., 2014; Hashimoto et al., 1995; Kadota et al., 2001; Kreis et al., 1993; van der Knaap et al., 1990) and a smaller longitudinal study (Holmes et al., 2017), providing strong evidence that tNAA levels rise across childhood within individuals. Increases in tNAA may be related to several aspects of fiber tract development as well as changes in metabolic activity. NAA is highly concentrated in neuron bodies and axons, so increasing levels of tNAA across childhood may indicate axonal development including elongation, increasing axon diameter, and/or increasing axon packing (Blüml et al., 2013; Ross & Sachdev, 2004). This interpretation is consistent with evidence from diffusion MRI studies, which show increases in axon packing and fiber cross-section across childhood (Dimond et al., 2020; Genc et al., 2017; Mah et al., 2017). Developmental increases in tNAA could also reflect myelination, given that NAA is present at substantial levels in myelin and oligodendrocyte cell bodies and plays a role in the synthesis of myelin (Nordengen et al., 2015; Rae, 2014). In gray matter, increasing tNAA levels could reflect elevated metabolic activity (Rae, 2014) and/or intracortical myelination, which continues to develop through childhood and adolescence (Norbom et al., 2020). In addition, NAAG may account for some of the observed increase in tNAA, and could mark metabolic activity and availability for synthesis of glutamate (Rae, 2014). Baslow (2015) proposed a role of NAAG metabolism in complex task performance, so NAAG may increase as children engage in more cognitively demanding tasks (e.g., reading, math). This is especially relevant in the two regions investigated here, which are involved in executive functioning and language/reading processes that develop throughout childhood (Braver et al., 2021; Numssen et al., 2021; Richlan, 2012).

tCho levels decreased with age in both regions studied. We observed a curvilinear trajectory in the longitudinal sub-sample, such that tCho decreased most rapidly in early childhood and stabilized in middle childhood, reaching a minimum value just before 9 years. Prior studies have shown the most rapid tCho decreases in the first two years of life (Blüml et al., 2013; Degnan et al., 2014; Hashimoto et al., 1995; Kreis et al., 1993; Vigneron, 2006), coinciding with a period of rapid myelination (Deoni et al., 2011). Phosphorylcholine, a main constituent of the tCho signal, is incorporated into myelin sheath macromolecules during myelination (Blüml et al., 1999). This process may drive developmental decreases in tCho. Cross-sectional evidence has shown that tCho declines into early childhood, but our longitudinal analysis suggests that minimum tCho levels are reached several years later than previously thought (Blüml et al., 2013). Much like the rapid tCho changes in the first two years of life, decreasing tCho in childhood coincides with ongoing myelination in major white matter tracts and within the cortex (Lebel & Deoni, 2018; Norbom et al., 2020). The curvilinear pattern of decrease in our longitudinal sub-sample aligns with a slowing of myelination in childhood (Lebel & Deoni, 2018). Importantly, tCho effects are not limited to myelin, and tCho reflects turnover and content of membranes more broadly and may be a marker of cell density (Rae, 2014). Thus, tCho decreases in gray matter could reflect changes in cell density in addition to intracortical myelination.

We found a small age-related increase in tCr levels in the LTP region, with linear effects in both the full sample and longitudinal sub-sample. However, this effect did not remain significant when including SNR in the model; likely because the inclusion of SNR accounted for a substantial amount of age-related variance for this already small effect. An age-related increase in tCr is consistent with gradual age-related changes reported in prior studies (Blüml et al., 2013; Degnan et al., 2014; Girard et al., 2006; Holmes et al., 2017; Kreis et al., 1993). tCr is involved in maintaining energy homeostasis in the brain and is abundant in areas of high synaptic activity (Rae, 2014); increasing tCr across development may reflect greater energy demands as the brain matures (Blüml et al., 2013). tCr levels may also be related to the amount of local synaptic activity (Rae, 2014). Our result may be associated with age-related increases in LTP activity, consistent with a prior finding from our lab which showed increases in resting-state local activity and global connectivity across early childhood in a corresponding region (Long et al., 2017). Age-related effects in tCr levels raise an important methodological consideration: tCr has been used as an internal reference for calculating metabolite ratios under the assumption that tCr levels are consistent over time and between individuals. However, this assumption is increasingly shown to be incorrect, hence a preference for water referencing (Near et al., 2020; Rae, 2014). The age-related increase in tCr observed here confirm that tCr-referencing is unsuitable in children.

Our longitudinal sub-sample analysis revealed a significant nonlinear pattern of age-related changes in Glx, even though no significant linear relationships were found in the full sample for either voxel. In the longitudinal sample, Glx increased gradually in early childhood, peaked at age 6.9 years, and subsequently declined. Prior studies have shown age-related increases in Glx in infancy and childhood, but findings have been mixed with regard to the timing of Glx fluctuation (Blüml et al., 2013; Degnan et al., 2014; Holmes et al., 2017). Studies of adolescents and young adults have shown age-related decreases in Glu and Glx (Devito et al., 2007; Ghisleni et al., 2015; Hädel et al., 2013; Marsman et al., 2013; Perica et al., 2022; Raininko & Mattsson, 2010; Shimizu et al., 2017; Volk et al., 2019), indicating that the decline in Glx observed in the older range of our sample likely continues through adolescence and into adulthood. Our longitudinal data provide additional power over prior cross-sectional and smaller studies to detect nonlinear patterns of development based on change within individuals, and to predict the age at which Glx levels peak (6.9 years based on our fitted model). The individual fit lines shown in Figure 5 reveal variability in individual trajectories (**∩** and U shapes) and ages at which Glx levels peak, highlighting the inter-individual variability in development. These individual growth patterns are of interest for future work investigating metabolites development in relation to cognitive and behavioral traits. Notably, Glx measurements exhibit higher levels of interindividual variability and measurement error relative to other metabolites (Soreni et al., 2010), so replication of this finding in an independent longitudinal sample is needed.

Glx may reflect metabolism, excitatory neural activity/excitability, and neuronal synchronization (Rae, 2014; Rodriguez et al., 2013; Stagg et al., 2011), but the interpretation of developmental changes in Glx levels is complicated due to the multifaceted role of glutamate as a metabolite and neurotransmitter. As a result, simultaneous developmental processes can involve co-occurring increases and decreases in Glx. For example, increasing Glx related to increasing excitatory neural activity may be counterbalanced by decreasing Glx related to pruning. The balance of these processes likely varies over time and across brain regions, contributing to interindividual variability in Glx measurements. The contribution of glutamine to the MRS signal further complicates matters, but the isolated Gln signal was not sufficiently reliable to facilitate examination of independent patterns of Glu and Gln development in our study. This should be a target of future research given that opposite directions of change in Glu and Gln have been observed across adulthood (Hädel et al., 2013) and in a rat model of infancy-early childhood development (Ramu et al., 2016). Glu and Gln are also involved in the synthesis of GABA (Rae, 2014), and could thus be associated with developmental changes in GABA (Porges et al., 2021).

Our finding of initial increase and later decrease in Glx may reflect tuning of excitability in neural networks that support cognition and language. An optimal balance of excitatory and inhibitory activity is needed for efficient functioning (Ferguson & Gao, 2018). Hyperexcitability due to imbalance in the glutamatergic system has been proposed as a source of impairment in developmental disorders including autism spectrum disorder (Rubenstein, 2010) and reading disorder (Hancock et al., 2017). In contrast, over-inhibition has been associated with stress and affective disorders (Page & Coutellier, 2019). The initial phase of increasing Glx observed here coincides with our previous findings of increasing activity and connectivity of the LTP from ages 2-6 years in an overlapping sample of participants (Long et al., 2017). The decline in Glx over the later ages of our sample (7-12 years) may reflect a phase of balancing excitatory and inhibitory activity through pruning of excitatory glutamatergic synapses (Lieberman et al., 2019) and/or increasing inhibitory GABAergic activity (Porges et al., 2021). Our data likely captures only the beginning of this phase; recent evidence shows declining Glu/Cr and increasing balance of Glu/Cr and GABA/Cr from adolescence into early adulthood (Perica et al., 2022).

We did not find significant changes in mI across the age-range studied, though we observed a trending decrease in the ACC dataset (p=.083). Prior cross-sectional studies show a decline in mI from infancy to adulthood (Blüml et al., 2013; Degnan et al., 2014; Kreis et al., 1993, 2002), with effects likely driven by the rapid changes that occur *in utero* and during infancy (Girard et al., 2006; Kreis et al., 1993). mI concentration varies regionally across the brain and is higher in gray matter than white matter (Rae, 2014), which may explain the presence of an age-related trend in the ACC, but not the LTP in our study. Further research is needed to test for region-and tissue-specific changes in mI from infancy through childhood.

We did not find any sex effects or interactions in our models, consistent with a previous longitudinal study in an overlapping age range (Holmes et al., 2017). Nonetheless, the null findings do not provide conclusive evidence for a lack of sex differences in metabolite development and future studies should continue to examine possible sex-related effects, especially in adolescent and adult samples, as such effects may emerge later in development.

### Limitations

Our study was limited by several methodological constraints. Metabolite levels are known to differ between gray matter and white matter (Rae, 2014), and differences in developmental patterns have been reported based on tissue type (Blüml et al., 2013). We corrected for cerebrospinal fluid and tissue specific relaxation effects in the calculation of metabolite concentrations according to recommended best practice (Gasparovic et al., 2006; Near et al., 2020). Gray matter tissue fraction was included as a covariate in our statistical models for the LTP data to control for individual differences in the tissue content of the voxels (tissue fraction improved model fit for the LTP data, but not for the ACC data, as the ACC voxels consisted almost exclusively of gray matter). Nonetheless, we cannot draw conclusions about different patterns of metabolite development in gray and white matter. In addition, the ACC and LTP datasets differed in numbers of participants, numbers of longitudinal timepoints, and ages at data collection, so the region-specific effects we report require replication in matched datasets. Further, we cannot generalize our findings to other regions of the brain. Data quality is a concern in young participants, and differences in data quality across development may have impacted results. We made an effort to ensure best data quality based on most recent recommendations for data processing and analysis, including retrospective frequency and phase correction and analysis with a customized basis set (Near et al., 2020; Simpson et al., 2017), along with visual inspection of all fitted spectra to confirm quality. Post-hoc analysis including quality metrics (SNR and linewidth) indicate that effects of data quality on our results were minimal.

### Conclusion

This study provides novel insight to the development of brain metabolites across early-middle childhood, building on prior work that has mainly focused on infancy. We found significant and substantial age-related effects in levels of tNAA and tCho across early-middle childhood in the ACC and LTP. Additional regional effects were found for tCr and Glx in the LTP. In addition to the overall model fits, our longitudinal data reveal inter-individual variability in the directions of change and ages of inflection points in metabolite development. These developmental effects parallel the structural and functional brain development of childhood (Faghiri et al., 2018; Frangou et al., 2021; Lebel & Deoni, 2018; Long et al., 2017; Norbom et al., 2021; Reynolds et al., 2019). Future research that integrates other imaging modalities will help to elucidate the mechanisms underlying these developmental changes. This knowledge can further be linked to cognitive, behavioral, and clinical traits to trace the roots of disorders and diseases that emerge in childhood and provide new directions for intervention and treatment.

## Acknowledgements

This work was supported by the Alberta Children’s Hospital Research Institute, and the Canadian Institutes of Health Research (IHD-134090, MOP-136797). Salary support was provided by the Canada Research Chair Program (ADH, CL), the Hotchkiss Brain Institute (MMD), the Cumming School of Medicine (MP, MMD), and the Killam Trusts (MP).

