## Supplementary Materials for "Changes in brain metabolite levels across childhood"

### Supplementary Analyses of Data Quality Metrics

**Table S1.** Summaries of models testing relationships between QC metrics and age in the ACC dataset. No significant age effects were found.

| | ACC - SNR | | | | | ACC - Linewidth | | | | |
|---|---|---|---|---|---|---|---|---|---|---|
| *Predictors* | *Estimates* | *std. Beta* | *CI* | *standardized CI* | *p* | *Estimates* | *std. Beta* | *CI* | *standardized CI* | *p* |
| (Intercept) | 59.04 | -0.04 | 51.51 – 66.56 | -0.23 – 0.15 | **<0.001** | 6.79 | 0.07 | 4.64 – 8.94 | -0.13 – 0.27 | **<0.001** |
| Age (years) | -0.52 | -0.05 | -2.29 – 1.24 | -0.24 – 0.13 | 0.559 | 0.20 | 0.07 | -0.30 – 0.70 | -0.11 – 0.25 | 0.433 |
| N | 101 $_{\text{subj\_id}}$ | | | | | 101 $_{\text{subj\_id}}$ | | | | |
| Observations | 112 | | | | | 112 | | | | |

**Table S2.** Summaries of models testing relationships between QC metrics and age in the LTP dataset. Age was significantly associated with the SNR of the NAA peak ($p$<.001).

| | LTP - SNR | | | | | LTP - Linewidth | | | | |
|---|---|---|---|---|---|---|---|---|---|---|
| *Predictors* | *Estimates* | *std. Beta* | *CI* | *standardized CI* | *p* | *Estimates* | *std. Beta* | *CI* | *standardized CI* | *p* |
| (Intercept) | 59.37 | 0.04 | 54.57 – 64.17 | -0.10 – 0.17 | **<0.001** | 7.14 | -0.03 | 6.33 – 7.94 | -0.16 – 0.10 | **<0.001** |
| Age (years) | 2.13 | 0.30 | 1.39 – 2.87 | 0.19 – 0.40 | **<0.001** | -0.09 | -0.08 | -0.22 – 0.04 | -0.19 – 0.03 | 0.157 |
| N | 95 $_{\text{subj\_id}}$ | | | | | 95 $_{\text{subj\_id}}$ | | | | |
| Observations | 318 | | | | | 318 | | | | |

**Figure S1.** Relationship between age and SNR of the NAA peak in the LTP dataset. Model fit lines shown in black; individual participants’ fit lines shown in red and blue for participants with multiple scans.

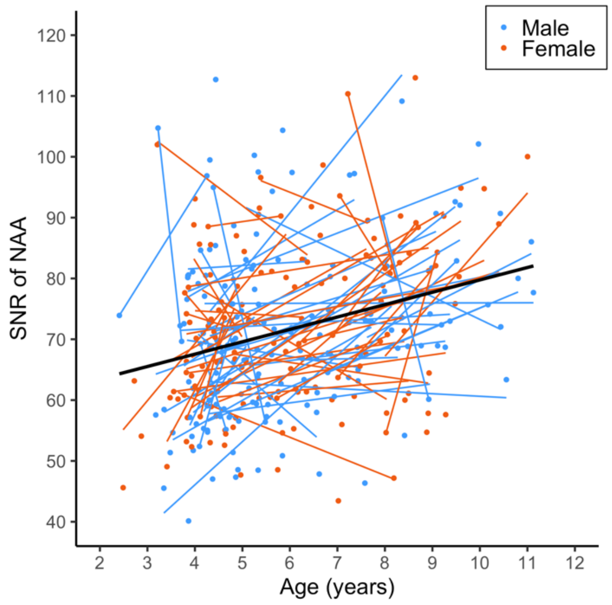

**Table S3.** Summary of age effects in the LTP, accounting for GM tissue fraction and SNR of the NAA peak. Significant p-values are presented in bold.

| | | | Age effect | | |
|---|---|---|---|---|---|
| **Metabolite** | **N obs.** | **Intercept** | **beta[95% CI]** | **Std. beta[95% CI]** | $p_{unc.}$ |
| tNAA | 317 | 10.69 | .16[.10, .22] | .29[.18, .40] | **<.001*** |
| tCho | 318 | 3.23 | -.04[-.06, -.03] | -.32[-.43, -.21] | **<.001*** |
| tCr | 318 | 6.43 | .04[-.01, .09] | .08[-.02, .18] | .118 |
| Glx | 317 | 14.13 | .07[-.04, .17] | .06[-.03, .15] | .205 |
| mI | 314 | 5.23 | -.01[-.08, .07] | -.01[-.14, .11] | .825 |
